# Are Different Populations Fairly Represented in Single-Cell Omic Atlases?

**DOI:** 10.1101/2025.10.01.677375

**Authors:** Catrina Yang, Kavitharini Saravanan, Aryan Saharan, Kuan-lin Huang

## Abstract

Single-cell Omic atlases are transforming biology and medicine, yet their demographic representativeness has not been systematically evaluated. We analyzed >13,500 samples from the Human Cell Atlas (HCA), Human Tumor Atlas Network (HTAN), and PsychAD Consortium. Benchmarking against global and US general and disease-prevalence data, we found a striking, pervasive European over-representation and under-representation of Asian and Latino individuals. Nearly 70% of HCA samples lacked ancestry annotation, and among annotated samples, Europeans were over-represented sixfold. PsychAD was nearly two-thirds European. HTAN tumors were 69% European, with several cancer types showing sex skews beyond expected incidence. These disparities highlight that current single-cell resources risk embedding inequities into AI foundational models, biomarker discovery, and therapeutic development. We contextualize these findings within the structural, economic, and regulatory, survey the growing ecosystem of diversity-focused initiatives, and provide an actionable, field-specific checklist to help research teams design single-cell studies whose benefits extend equitably across populations.

## MAIN

Single-cell omics technologies have revolutionized our understanding of cellular heterogeneity, enabling the discovery of rare cell types, transitional states, and microenvironmental dynamics that bulk approaches obscure. Large-scale initiatives such as the Human Cell Atlas (HCA)^1-3^, the Human Tumor Atlas Network (HTAN)^4,5^, and the PsychAD consortium^4,6^ have generated expansive datasets spanning healthy tissues, tumor microenvironments, and neuropsychiatric or neurodegenerative conditions. These resources are invaluable to biomedical discovery, but systematic evaluation of their demographic composition has been limited, raising concerns about their representativeness and generalizability.

The challenge of representation has long plagued human genomics. Genome-wide association studies (GWAS) have been overwhelmingly conducted in individuals of European ancestry, leading to biased polygenic risk scores (PRS) that transfer poorly to other populations^7,8^. PRS derived from European cohorts often lose their predictive accuracy when applied to non-European groups^9,10^. Beyond causing technical issues for research, the lack of diversity in genomics risks reinforcing health disparities when biomedically-actionable findings are disproportionately relevant to majority populations^11^.

Genetic ancestry shapes cell-type-specific gene expression and regulatory architecture. Single-cell studies of viral immune responses have identified hundreds of genes with ancestry-differential expression in specific immune cell subsets^12,13^, and patient ancestry modulates innate immune activation at the level of individual cell states in SARS-CoV-2 responses^14^. An atlas from one ancestral background, therefore risks missing cell subtypes or wrongly assuming universal cell markers—paralleling how the original reference genome missed variants or genome segments now recovered by the Human Pangenome Project^15^. While definitive population-specific cell types or cell states remain to be proven, many pathogenic variants are unique to specific populations^16,17^, and ancestry-driven regulatory variation can shift cell states in disease-relevant cell populations that a homogeneous atlas would fail to catalog. Diverse single-cell maps are therefore a prerequisite for biological completeness—a *pancellular* reference that, like the pangenome, must represent humanity’s full diversity to be scientifically universal^18,19^.

Single-cell consortia, where significant resources are unified to survey cellular profiles in thousands of individuals, risk reproducing inequities in genomics. The outputs of these large-scale studies are often marketed as universal reference maps of human biology and used to train foundation models^2,20-22^. However, recent studies have suggested that the profiled cellular omic signatures could indeed be ancestry- and sex-specific^18,19^, calling for the need to establish single-cell datasets that fairly represent different populations. Here, we systematically evaluate ethnicity/race and sex representation of sampled individuals in three major single-cell consortia-generated atlases (**Table 1**), providing a diversity audit to inform future study designs.

**Table 1.**
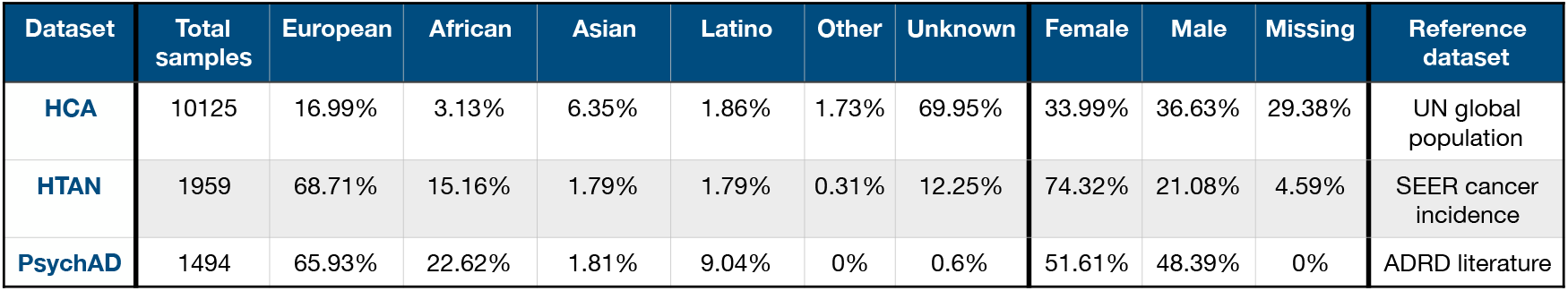
Overall ancestry and sex composition of individuals from the Human Cell Atlas (HCA), Human Tumor Atlas Network (HTAN), and the PsychAD consortia evaluated in this study.

### Human Cell Atlas (HCA) vs. Global Population Demographics

The HCA dataset comprised 10,125 samples across 15 tissue categories^1,2^, the largest included immune (n = 3,704, 36.6%), gut (n = 883, 8.7%), skin (n = 906, 8.9%), lung (n = 743, 7.3%), and kidney (n = 601, 5.9%)(**Supplementary Figure 1, Supplementary Table 1**). Overall, 69.9% of samples lacked ancestry annotation after we thoroughly cataloged data from the HCA website or Supplementary Information reported by the linked publications (**Methods**). Among the 3,043 samples with reported ancestry, Europeans comprised 56.5%, followed by Asians (21.1%), Africans (10.4%), Latinos (6.2%), and Other (5.8%)(**Figure 1A**). Compared to global expectations (Europe 9.4%, Asia 59.4%, Africa 17.6%, Latino 13.6%), Europeans were overrepresented sixfold, while Asians, Africans, and Latinos were consistently underrepresented (0.36×, 0.59×, and 0.45×, respectively; p < 2e-16). We acknowledge that the use of the global reference is an oversimplification and provides only coarse categories that align across studies after the extensive curation of a wide range of ancestry metadata (**Methods**).

**Figure 1.**
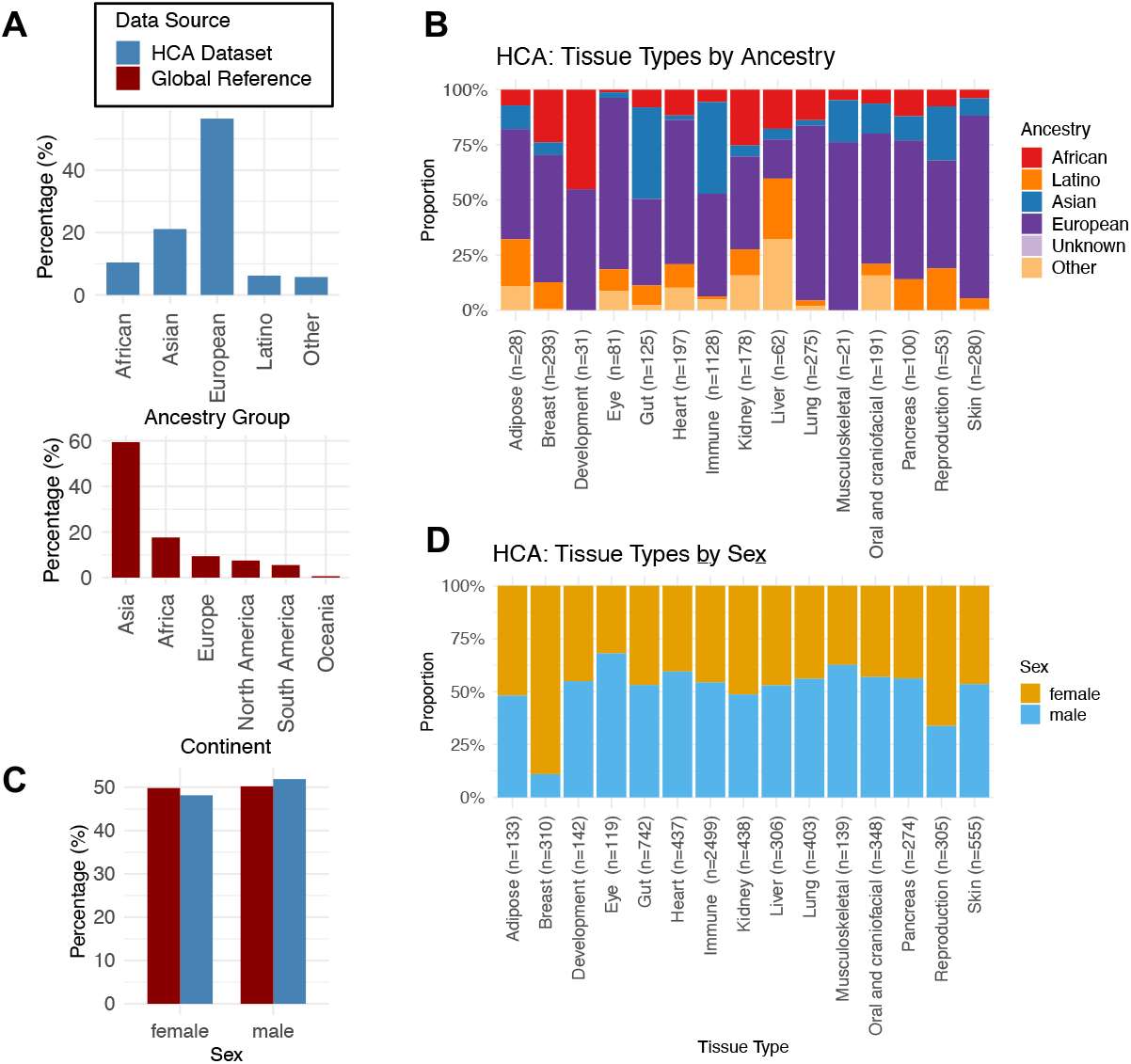
Human Cell Atlas (HCA) ancestry and sex composition across tissues. (A) Comparison of ancestry proportions in HCA (excluding samples with unknown ancestries, in red bars), with proportions of European, African, Asian, Latino, Other ancestry to the global reference data of populations across continents (in blue bars). (B) Stacked barplots of ancestry distributions across 15 HCA tissue categories (excluding samples with unknown ancestries), with proportions of European, African, Asian, Latino, Other, and Unknown ancestry shown for each tissue. (C) Comparison of sex proportions in HCA (excluding samples with unknown sexes, in red bars) to global population reference distributions (in blue bars). (D) Stacked barplots of sex distributions across 15 HCA tissue categories (excluding samples with unknown ancestries), with proportions of female and male shown for each tissue.

Because conclusions drawn from complete-case analysis alone could be severely biased if missingness is non-random, we considered a simple thought experiment: how would the observed European share change at two extremes: if every unlabeled donor turned out to be European (pushing the share to its highest possible value, or ceiling), versus if none of them were European (pushing it to its lowest possible value, or floor)? The true share has to lie somewhere between these two extremes, regardless of why annotations are missing. We refer to this approach as partial-identification bounds, where the upper bound (ceiling) assumes all missing donors belong to the group of interest and the lower bound (floor) assumes none do (**Supplementary Table 2**). While other ancestries’ underrepresentation becomes less stable under conservative missingness estimates, European over-representation is robust to all plausible missing-data mechanisms: even under the most conservative assumption (all 7,082 missing donors are non-European), the European lower bound (17.0%) still exceeds the global reference (9.4%).

**Table 2.** Practical Checklist for Equitable Single-Cell Omic Study Design.

| # | KEY QUESTION FOR RESEARCH TEAMS | POSSIBLE SOLUTIONS |
| --- | --- | --- |
| **Recruitment & Study Design** |  |  |
| 1 | Does the planned cohort reflect population incidence/prevalence for the target tissue/disease? | • Set explicit sampling quotas by ancestry<br>• Oversample underrepresented groups |
| 2 | Is sex balance planned for non-sex-specific tissues/diseases? | • Set sex-stratified quotas |
| 3 | Have we defined minimum per-group sample sizes to enable stratified analyses by ancestry and/or sex? | • Power analyses by subgroup; enforce thresholds<br>• Phase enrollment until quotas are met |
| 4 | Are community sites serving underrepresented populations included as recruitment hubs? | • Add community clinics & advisory boards<br>• Culturally tailored outreach & consent |
| 5 | Are eligibility criteria free of inadvertent exclusions (language, insurance, visit schedule)? | • Translated materials & flexible visit windows<br>• Reimbursements/transport, childcare support |
| **Pre-Analytical & Single-Cell-Specific** |  |  |
| 6 | Do tissue procurement and dissociation protocols account for donor availability constraints across sites (e.g., organ-donor programs, surgical schedules)? | • Pre-qualify multiple procurement pipelines per site<br>• Standardize ischemia time, dissociation enzyme/duration, and viability thresholds |
| 7 | Are biobanking SOPs harmonized across sites (time-to-freeze, buffers, RNase control)? | • Unified SOPs with site audits and QC dashboards<br>• Reject out-of-spec samples uniformly |
| 8 | Do multiplexing and batch-assignment strategies avoid confounding ancestry/sex with technical batch? | • Randomize or block donors across pools/batches by ancestry and sex<br>• Include cross-site reference samples in every run<br>• Use equitable barcoding allocation (balance hashtag/lipid oligos across groups) |
| **Metadata & Monitoring** |  |  |
| 9 | Will ancestry, sex, age, and disease-specific clinical variables be mandatory fields using controlled vocabularies? | • Require fields at upload using ontology standards (e.g., cellxgene schema, EBI SCEA metadata)<br>• Validate against HAncestro for ancestry |
| 10 | Are missingness dashboards (ancestry × sex × site) reviewed regularly? | • Publish dashboards; open action items to sites when gaps emerge |
| 11 | Is there a plan for *post-hoc* enrichment if monitoring shows imbalance? | • Pre-specify trigger rules (e.g., <20% of target quota)<br>• Add contingency recruitment sites; targeted outreach |
| 12 | Will raw + harmonized metadata be versioned and publicly released? | • Robust versioning & changelog on every data release |
| **Community Engagement & Ethics** |  |  |
| 13 | Have we planned return-of-value to sustain participation from diverse communities? | • Share lay summaries and participant newsletters<br>• Host community feedback sessions |
| 14 | Does consent language address data sovereignty, international sharing, and re-use in AI model training? | • Consult local governance frameworks (e.g., CARE Principles for Indigenous data)<br>• Include explicit clauses on computational re-use and AI model training |
| **Downstream AI/ML Model Development** |  |  |
| 15 | Will training-data demographic composition be disclosed in model documentation? | • Report ancestry/sex breakdown of all training and test splits<br>• Flag known gaps relative to target application population |
| 16 | Will model performance be evaluated and balanced when stratified by ancestry and sex? | • Report cell-type annotation accuracy, embedding quality, and cell marker predictions per demographic group<br>• Include robustness tests under simulated demographic shift |

Tissue-level analyses revealed pronounced but heterogeneous skews—all tests compare observed ancestry mix within tissue to the global reference mix (see **Methods**) showed FDR <0.01, except for musculoskeletal where 205/226 were of unknown ancestry (**Figure 1B**). Among ancestry-labeled samples most biased for European samples, skin was 82.9% European, lung 79.3%, eye 77.8%, musculoskeletal 76.2%, and heart 65.5% European. By contrast, immune and gut showed comparatively higher Asian representation (41.7% and 41.6%, respectively), with lower European fractions (immune 46.5%, gut 39.2%). African ancestry individuals were better represented in development (45.2%), kidney (25.3%), and breast (23.9%) atlases, whereas Latino individuals showed higher representation in liver (27.4%), adipose (21.4%), and reproduction (18.9%) atlases (**Figure 1B**). These skews in ancestral proportions likely reflect study-purpose-driven sampling rather than intentional exclusion: many HCA datasets derive from tissue-banking programs with non-random recruitment (organ donors, surgical excess, pediatric samples) whose demographics reflect the population served by contributing institutions. We note that high ancestry-missingness in HCA (69.9%) likely obscures the true ancestry compositions and should be addressed in future releases.

Sex ratios in HCA were 34.0% female, 36.6% male, and 29.4% unreported (**Supplementary Figure 1**)(**Figure 1C**). Among samples with sex information, expected skews were observed in breast (89.0% female) and reproductive tissues (66.2% female). Several tissues were modestly male-enriched, including immune (54.3% male), eye (68.1% male), lung (56.1% male), and musculoskeletal. No significant sex biases were observed for adipose, development, gut, kidney, and liver (P>0.17), suggesting more balanced sampling in these tissue atlases (**Figure 1D**). These sex patterns are sometimes directionally consistent with physiological expectations in sex-specific disease tissues, but also suggest potential ascertainment or study-design bias elsewhere.

### The Human Tumor Atlas Network (HTAN) vs. the US Cancer Population Incidence

The HTAN dataset comprised 1,959 tumor samples across 22 cancer types ^4,5^. Breast cancer was the largest contributor (n = 993, 50.7%), followed by lung (n = 245, 12.5%), colorectal/colon (n = 151, 7.7%), skin (n = 79, 4.0%), neuroblastoma (n = 74, 3.8%), pancreas (n = 47, 2.4%), ovary (n = 26, 1.3%), and several smaller categories (<20 cases each)(**Supplementary Figure 2, Supplementary Table 3**). Overall, 12.3% of samples lacked ancestry annotation. Among the 1,719 samples with reported ancestry, Europeans comprised 78.3%, Africans 17.3%, Asians 2.0%, Latinos 2.0%, and Other 0.3%.

**Figure 2.**
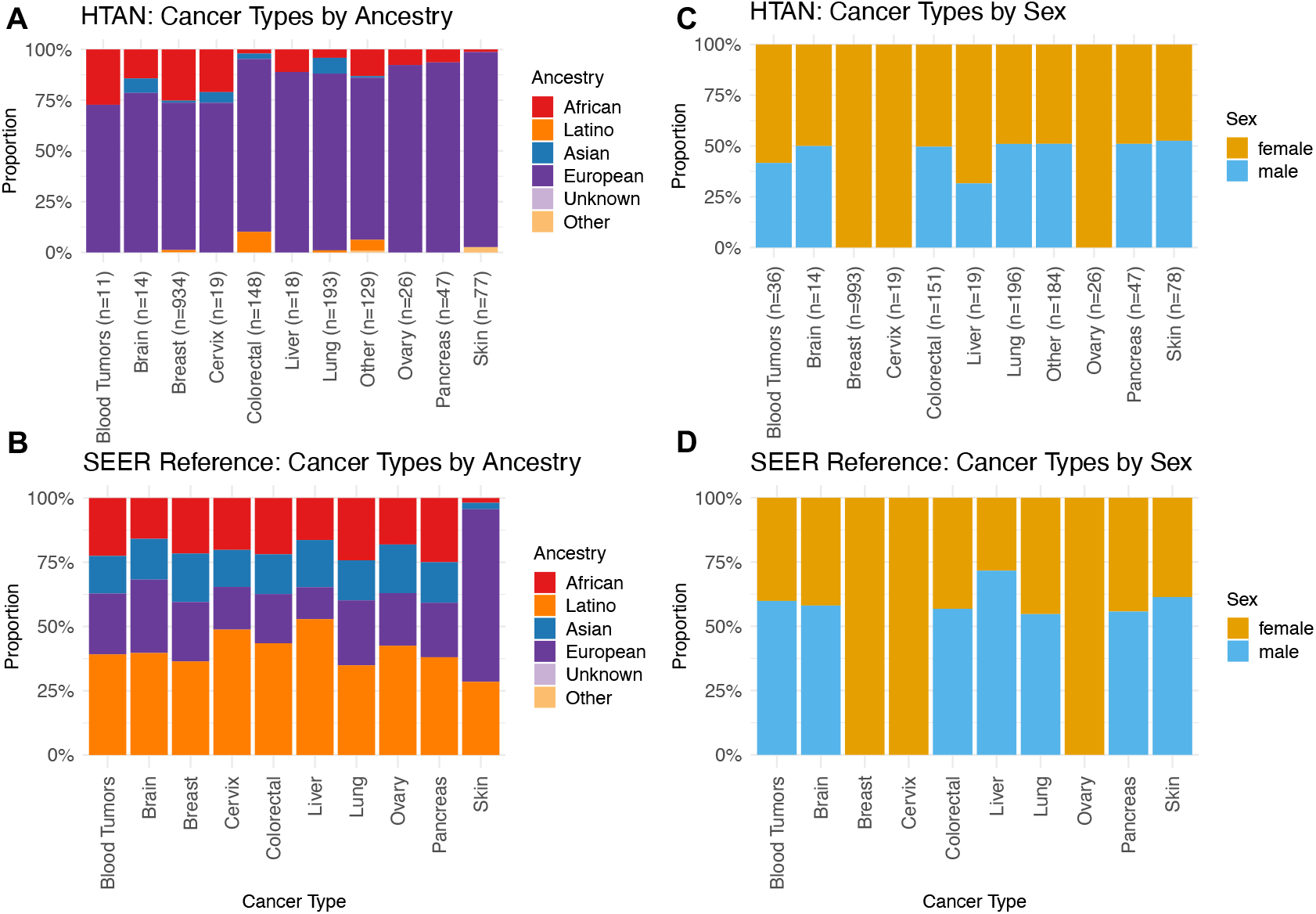
Human Tumor Atlas Network (HTAN) ancestry and sex composition across cancer types. (A-B) Stacked barplots comparing ancestry distributions across 11 major cancer categories in (A) HTAN (excluding samples with unknown ancestry) vs. (B) the reference SEER cancer incidence data of the respective cancer type, as shown in proportions of European, African, Asian, Latino, Other, and Unknown ancestry. (C-D) Stacked barplots comparing sex proportions in (C) HTAN (excluding samples with unknown sex) to (D) the reference SEER cancer incidence data of the respective cancer type.

Cancer-specific ancestry distributions revealed additional skew in several cancer types compared to their expected US incidences (**Figure 2A-B**). Within samples with annotated ancestry data, all the cancer tissue sites showed over 72% European. For example, skin cancer showed 96.1% European, pancreas 93.6% European, ovary 92.3% European, liver 88.9% European, and lung 87.1% European. Substantial African representations were found in blood tumors (27.3%), breast (25.3%), cervix (21.1%), and neuroblastoma (17.8%). Asian and Latino participants were severely under-represented in all cancer tissue sites; the highest count of Asian was in lung (N=15/245), which comprised 7.8% of the cohort, whereas the highest count of Latino was found in colorectal cancer (N=15/151)(**Figure 2A-B**).

The overall sex distribution was heavily female (74.3% female, 21.1% male, 4.6% unreported)(**Supplementary Figure 2**), driven by the predominance of breast cancer that comprised over half the cohort and minorly by other tissue sites, i.e., ovary, cervix, that were female-specific. Considering only cases with reported sex, multiple cancer types showed a female skew in HTAN, such as liver (68.4% female), lung (49%), blood (58.3%), and skin (47.4%) tumors, compared to the SEER female/male incidence of those respective cancers (**Figure 2C-D**). These sex patterns are sometimes directionally consistent with physiological expectations in sex-specific disease tissues, but also suggest potential ascertainment or study-design bias elsewhere.

### The PsychAD Consortium vs. US Demographics in Neurological Diseases

The PsychAD dataset comprised 1,494 samples drawn from three contributing cohorts: HBCC (n = 300), MSBB (n = 1,042), and RADC (n = 152)^4,6^. Across the cohort, diagnostic categories included Alzheimer’s disease (AD_combined, n = 373, 25.0%), schizophrenia (SCZ, n = 120, 8.0%), dementia with Lewy bodies (DLBD, n = 112, 7.5%), vascular dementia (n = 85, 5.7%), bipolar disorder (BD, n = 55, 3.7%), tauopathies (n = 47, 3.1%), Parkinson’s disease (PD, n = 40, 2.7%), frontotemporal dementia (FTD, n = 15, 1.0%), and dementia not otherwise specified (n = 439, 29.4%); in PsychAD, some individuals may have multiple diagnosis and 597 individuals who do not have any of these diagnoses (**Supplementary Figure 3, Supplementary Table 4**).

**Figure 3.**
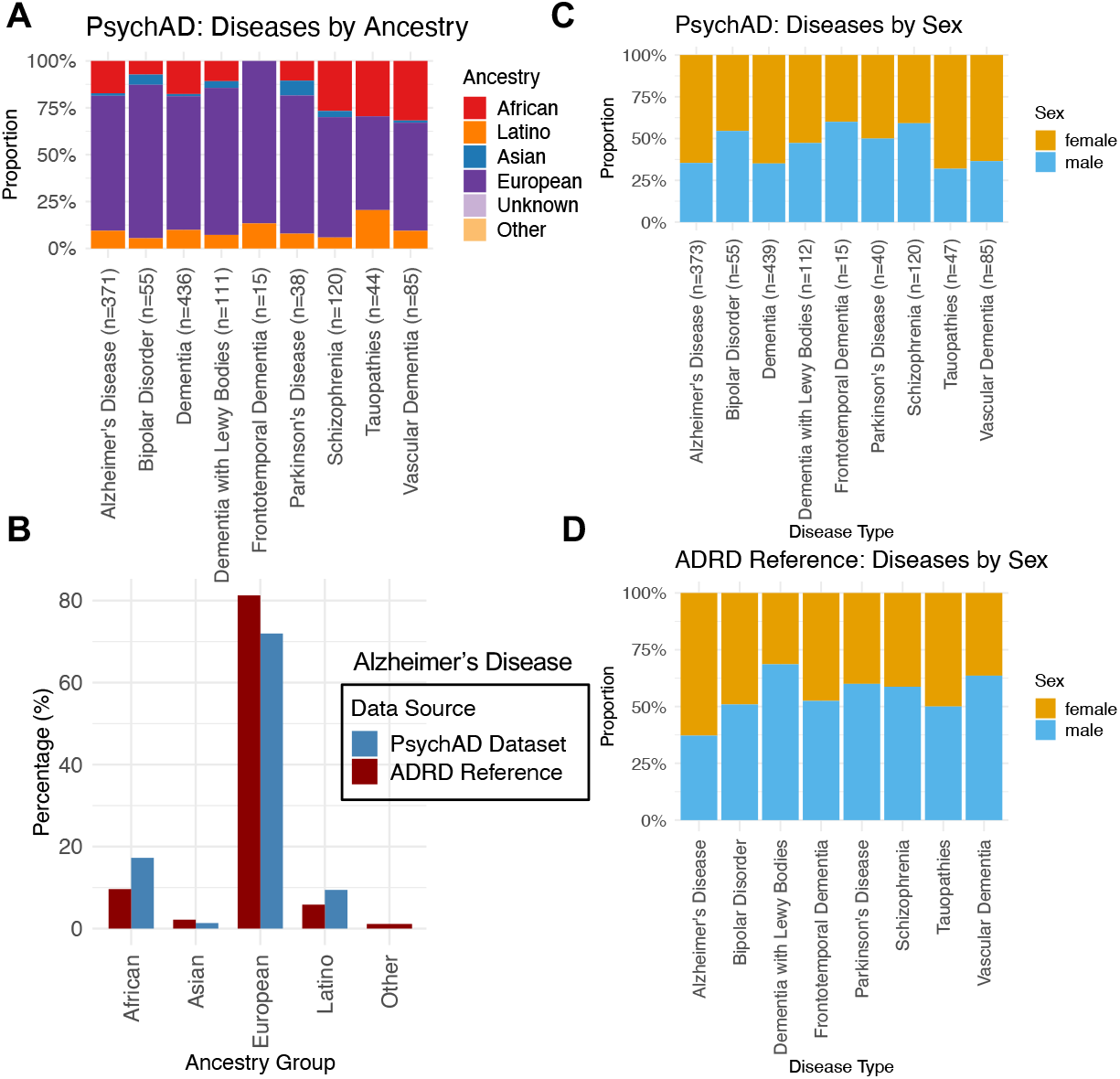
PsychAD ancestry and sex composition across neuropsychiatric and neurodegenerative diagnoses. (A) Stacked barplots of ancestry distributions across 9 diagnostic categories, including Alzheimer’s disease, schizophrenia, dementia with Lewy bodies, vascular dementia, bipolar disorder, tauopathies, Parkinson’s disease, frontotemporal dementia, and dementia not otherwise specified. (B) Comparison of ancestry proportions in PyschAD Alzheimer’s disease cases (excluding samples with unknown ancestries, in red bars), with proportions of European, African, Asian, Latino, Other ancestry to the reference data of US Alzheimer’s disease and related dementia cases (in blue bars). (C-D) Stacked barplots comparing sex proportions in (C) PyschAD (excluding samples with unknown ancestries) to (D) the reference data from literature for each dementia type.

Among all 1,494 Psych samples, Europeans accounted for 65.9%, Africans 22.6%, Latinos 9.0%, Asians 1.8%, and Unknown 0.6%. Compared to ADRD reference demographics in the United States (Europeans 59.1%, Africans 12.2%, Latinos 9.2%, Asians 6.6%)(**Methods**) (**Figure 3A-B**), AD cases in PsychAD were characterized by overrepresentation of Europeans (71.6%), slightly higher representation of Africans (17.2%) and near parity Latino (9.4%), but marked under-representation of Asians (1.4%). For other diseases, Europeans are heavily represented in FTD (86.7%), BD (81.8%), and DLBD (77.7%). African individuals comprised a considerable fraction of Vascular dementia (31.8%) and SCZ cases (26.7%). The highest percentage of Latino individuals were found within Tau (19.2%) and FTD (13.3%) cases. The highest fraction of Asian was found in PD and BD with mere 7.5% and 5.5%, respectively (**Figure 3A-B**).

Overall sex balance was near parity: 51.6% female and 48.4% male (**Supplementary Figure 3**). Disease-level analyses revealed stronger deviations from equal representation, and disease-specific sex ratios was compared to their respective reference data (**Methods**) (**Figure 3C-D**). AD and dementia not otherwise specified were 64.6/64.9% female, consistent with known female predominance. Vascular dementia also showed female enrichment in PyschAD (63.5% female). On the other hand, male skews were seen in SCZ (59.2% male, aligned with literature’s 58.7%), BD (54.6%), FTD (60%), and control (57.8%). While PD had 20 males and 20 females in PyschAD, the known disease incidence is about 60% male (**Figure 3C-D**).

### Significant Ancestry and Sex-Biases Exist in Surveyed Single-Cell Atlases

Our systematic assessment across HCA^1,2^, HTAN^4,5^, and PsychAD^4,6^ shows that demographic bias is pervasive in single-cell atlases. HCA had nearly 70% missing ancestry annotation, and among labeled donors, European individuals dominated all tissue types. HTAN revealed even starker distortions: Europeans comprised >78% of annotated samples and were over-represented in every cancer type compared to SEER US cancer incidence, while Asians and Latinos were severely under-represented in all cancer types. PsychAD was nearly two-thirds Europeans, with African ancestry better represented at 22.6% (higher than expected based on ADRD prevalence), but Asians were again underrepresented (1.8%). Sex balance was dataset-dependent: near parity in PsychAD overall, male skewing in multiple HCA immune tissues, and female skewing in HTAN across several non–sex-specific cancers.

These findings extend the well-documented inequities of genomics^7,9,11^. Just as GWAS and PRS suffer from limited transferability across ancestries^8,10^, single-cell resources biased toward European donors risk producing cell and disease maps that fail to generalize globally. This is not a trivial concern: genetic background influences gene expression and regulatory variation^23^, sex differences shape immune responses and disease trajectories^24^, and ancestry-associated biology can alter therapeutic options and responses, e.g., in genetics, *CYP2D6* allele frequencies governing metabolism of ∼25% of clinical drugs differ markedly across populations^25,26^. While single-cell signatures have yet been widely used for clinical stratification, over-representation of Europeans in single-cell genomics precludes discovery of biology that exists across the wide spectrum of human diversity and risks building single-cell models that obscure the cellular mechanisms most relevant to underrepresented groups.

This study has several limitations. First, ancestry was analyzed as reported metadata rather than genetically inferred, which may introduce misclassification. Classifying ancestry groups into broad categories like “European” or “Asian” is challenging due to the complex and personal nature of ethnic identity. These broad categories collapse substantial within-group heterogeneity and conflate self-identified race/ethnicity with genetic ancestry. Where metadata permitted, we sought finer granularity (*see Methods: Curation of Demographic Data in Single-Cell Consortia Studies*), but most datasets did not support it. Some groups, such as Ashkenazi Jews, classified as “European,” have distinct cultural and genetic backgrounds that may not fit neatly into this category. Similarly, “Hispanic/Latino” encompasses individuals with mixed heritage. Second, the high degree of missing data limits resolution, and small sample sizes for certain groups reduced statistical power for some disease-specific comparisons. Third, our analyses of ancestral/sex proportions do not resolve whether the imbalances are a result of study-purpose-driven sampling or intentional/unintentional exclusion. Fourth, the use of reference populations herein is limited by their surveyed populations and timepoints, for example, the global demographics of continental population shares used for HCA will not match genetic ancestry estimates. Finally, our analyses focused on 3 publicly available single-cell consortium datasets, which may not reflect the demographic composition of all other single-cell omic resources. Non-consortium datasets collectively have exceeded these consortia in sample size and may differ in composition, but often offer even less standardized metadata for analyses. Conclusions herein are based on data from these specific consortia, not the field as a whole. Despite these caveats, the consistency of European overrepresentation and Asian/Latino underrepresentation across these datasets underscores the robustness of our conclusions.

### How Can We Fix this Disparity?

The implications of disparity in single-cell omics are both scientific and societal. Scientifically, biased datasets constrain the interpretability of findings, hinder reproducibility, and limit the scope of discovery. Missing metadata compounds these problems, as incomplete annotation disproportionately affects careful study designs that account for diversity and limits the interpretability of cross-population analyses. Societally, they risk exacerbating health disparities by embedding inequities into the reference frameworks that guide biomarker discovery, drug development, and precision medicine. This can lead to challenges when a significant portion of future advances may be driven by AI foundation models built on these skewed datasets. For example, single-cell AI foundation models that are already widely used, such as scGPT^20^, Geneformer^27^, and scFoundation^28^ maximized the size of their training datasets by combining existing public datasets. Consequently, most foundation models include HCA as one of the major sources and leverage HTAN data if they separately trained a cancer-focused model without, to our knowledge, supplementation of data from underrepresented populations ^20-22^, thus possibly inheriting the bias from these datasets without most users’ awareness. A controlled benchmark of AI model performance stratified by donor ancestry remains needed.

Moreover, most AI tools are developed in Western institutions with GPU infrastructure; researchers in low- and middle-income countries may lack resources to run these models, creating a secondary inequity where analytical tools—not just reference data—reflect a narrow perspective. Providing technology-access programs in both single-cell and AI technologies is needed to help address such disparity.

The demographic skews emerge from intersecting structural forces. Most atlas projects draw samples from readily available sources at major institutions, often defaulting to local demographics unless diversity is considered during study design. Funding for large-scale projects has historically come from wealthier countries without explicit diversity requirements. Large research institutions that conduct such studies may also concentrate on certain geographical regions where recruited samples do not reflect the diversity of national or global populations. Single-cell technology remains expensive, limiting participation from low-resource settings. National data-governance policies create additional constraints: many countries restrict sharing individual-level genomic data outside their borders. Finally, achieving equity requires respectful community engagement—many populations have historical or cultural reasons for caution, and frameworks such as Indigenous data sovereignty principles^29^ appropriately constrain how data can be shared.

The research community is actively addressing these gaps. The HCA Equity Working Group and Diversity Task Force track and encourage donor diversity reporting^30^. In 2021, CZI launched Ancestry Networks for the HCA funding single-cell data collection from underrepresented populations worldwide, supporting projects including African, Latin American, and Asian immune diversity atlases and ancestrally inclusive organ atlases^31^. A recent framework from the HCA outlines governance, inclusive funding, and cross-regional partnership principles for equitable consortia^1^. Technically, initiatives to estimate genetic ancestry directly from scRNA-seq data via SNP-based approaches^32^ may provide ancestry estimates for nearly all atlas samples, superseding reliance on self-reported labels. Data portals, including cellxgene and EBI’s Single Cell Expression Atlas are incorporating structured ancestry fields in metadata schemas.

Moving forward, several structural reforms are still needed. First, intentional recruitment strategies should align donor sampling with disease incidence and population demographics^33^, as proposed in recent roadmaps for equity in genomics^9^. Second, metadata completeness and standardization should be mandated in single-cell repositories, with ancestry, sex, and tissue/disease-specific clinical variables treated as required fields. This can be achieved by harmonized toolsets and checklists such as those provided by the HCA Ethics Working Group^34^. Indeed, datasets such as PsychAD has much more complete metadata, suggesting this could be achieved with careful study and workflow design. Third, equity metrics should be adopted as benchmarks for evaluating single-cell datasets, complementing measures of resolution and scale with measures of demographic diversity.

To translate this into actionable steps, we developed a field-specific checklist (**Table 2**) covering 16 specific questions and 5 domains across the full study lifecycle: procurement and recruitment planning aligned to sampling frame; pre-analytical considerations including tissue dissociation and donor availability; equitable multiplexing and batch-assignment strategies to avoid confounding demographics with batch effects; minimum required demographic metadata fields and controlled vocabularies (aligned with cellxgene and EBI standards); standardized biobanking protocols; consent language and data-sovereignty considerations for international collaborations; and AI model-development reporting standards including training-data composition disclosure, stratified performance evaluation across ancestry and sex, and robustness testing. We include the AI/ML model-development items within this single-cell checklist because cellular reference atlases and the AI foundation models trained on them are now produced as a continuous pipeline: the same study-design choices that determine donor diversity at recruitment propagate directly into the training corpora of downstream cell-state classifiers and foundation models. Embedding these considerations within one checklist clarifies accountability. Each domain pairs a critical question with concrete solutions—for example, stratified recruitment quotas, community clinic partnerships, and pre-specified minimum sample sizes per ancestry group. This approach aims to address bias proactively, at the point of study design, where correction is most effective.

Single-cell omics risks repeating inequities in genomic studies—with consequences extending to the cellular reference maps that define human biology. Unless corrective actions are taken at the level of study design, consortium governance, funding policy, and technical infrastructure, the resulting benefits may become unevenly distributed. The HCA Equity Working Group, CZI Ancestry Networks, and regional initiatives represent real progress. Our analysis provides a quantitative baseline across three major consortia and provides a framework against which this progress can be measured. A *pancellular* reference representing humanity’s full ancestral, sex, and environmental diversity is the only atlas that will deliver on single-cell biology’s translational promise for all populations.

## METHODS

### Study Design and Data Sources

We conducted a systematic evaluation of sex- and ancestry-related representation across three large-scale single-cell resources: the Human Cell Atlas (HCA), the Human Tumor Atlas Network (HTAN), and the PsychAD consortium. These datasets were chosen because they represent flagship community efforts to generate, harmonize, and disseminate high-resolution cellular and disease-relevant data. For contextual benchmarking, we compared observed representation in each dataset against appropriate reference distributions: (1) global ancestry proportions from United Nations population statistics for HCA; (2) SEER cancer incidence rates stratified by ancestry and sex for HTAN; and (3) Alzheimer’s disease and related dementias (ADRD) prevalence estimates for PsychAD. Each of these data sources were described in detail in the later sections of the Methods.

In brief, metadata files from HCA, HTAN, and PsychAD were harmonized into canonical categories. Ancestry was standardized into African, Asian, European, Latino, Other, and Unknown. For HTAN, Hispanic/Latino identifiers were mapped to “Latino” regardless of race; for PsychAD, genetic ancestry labels (e.g., EUR, AFR, AMR, EAS, SAS) were collapsed into the canonical categories. Biological sex was standardized to female, male, or missing. For disease and tissue annotations, HCA samples were grouped by tissue of origin (15 categories), HTAN samples by cancer type (22 categories), and PsychAD samples by diagnosis (9 categories).

### Statistical Analyses

We computed descriptive statistics for ancestry and sex distributions overall and within tissues or disease categories. Representation was quantified both as raw counts and as proportions among non-missing annotations. Chi-square tests compared observed vs. expected representation, using population-level baselines (global ancestry, SEER, or ADRD). Multiple testing corrections were further conducted using the BH procedure for FDR. To measure magnitude of imbalances, we also calculated representation ratios (observed ÷ expected), equity indices (symmetrized measures of under-vs. overrepresentation), and diversity metrics (Simpson’s and Shannon’s indices) in the code-generated logs.

For HCA, analyses were stratified by tissue type; for HTAN, by cancer type; and for PsychAD, by diagnosis. Within each stratum, we assessed ancestry composition, sex distribution, and statistical significance relative to reference populations. We also quantified missingness for ancestry and sex. All analyses were performed in R using packages including *dplyr, tidyr, ggplot2, car*, and *scipy*. Scripts were version-controlled, and all transformations were logged for reproducibility.

### Global reference population

We used share of the world population by major world region (continent). Specifically, we used the widely-cited 2021 distribution: Asia 59.4%, Africa 17.6%, Europe 9.4%, North America 7.5%, South America 5.5%, Oceania 0.6%. These values correspond to UN-based tallies compiled in the “World population by continent, 2021” table^35^. Global sex composition was derived from the United Nations World Population Prospects (WPP) 2022 summary, which reports that the world had 50.3% males and 49.7% females in 2022^36^.

### U.S. Cancer reference population

We used SEER*Explorer (National Cancer Institute) to obtain age-adjusted incidence rates (per 100,000) for 2018–2022 by cancer site, race/ethnicity, and sex. SEER*Explorer methods specify that rates are age-adjusted to the 2000 U.S. standard population and are reported per 100,000 persons; incidence estimates incorporate SEER delay-adjustment methodology^37^.

For ancestry/ethnicity, we used the five mutually exclusive SEER race/ethnicity categories:

- Non-Hispanic White
- Non-Hispanic Black
- Non-Hispanic American Indian/Alaska Native (AI/AN)
- Non-Hispanic Asian/Pacific Islander (API)
- Hispanic (any race)

These are standard, mutually exclusive SEER groupings (Hispanic supersedes race). Sex was coded as Male and Female.

SEER provides rates rather than percentage shares. To construct reference distributions (for ancestry and for sex) within each *cancer site*:

1. We retrieved age-adjusted incidence rates for the target grouping (e.g., five race/ethnicity groups; or the two sex groups).
2. For each cancer site, we converted rates to shares by dividing each group’s rate by the sum of rates across groups for that site (i.e., compositional normalization so that shares sum to 1.0 within the site).

These normalized shares were used as the U.S. Cancer reference for ancestry/ethnicity and sex composition, respectively. Because rates are age-adjusted, the share is a proxy for the composition of incident cases, not general population composition.

### ADRD reference population

#### U.S. ancestry/ethnicity composition (ADRD)

For Alzheimer’s disease and related dementias (ADRD), we used the analysis by Matthews et al. (2019), which reported age-standardized prevalence estimates by race/ethnicity in adults aged ≥65 years^38^. Specifically, they reported the 2014 number of Medicare Fee-for-Service (FFS) beneficiaries with ADRD in each of the NH white, black, Asian and Pacific Islander (annotated as Asian for our analysis), Hispanic (annotated as Latino for our analysis), and American Indian and Alaska Native as well as Other (grouped into Others for our analysis). For the other diseases, we could not find consistent race-specific incidence data and thus were not modelled.

#### Sex composition

To standardize sex differences in incidence and prevalence across neuropsychiatric and neurodegenerative diseases in the United States, we relied on the best available population-based studies and large systematic reviews, converting reported sex-specific rates or ratios into fractions that sum to one. For Alzheimer’s disease and related dementias (AD/ADRD), we adopted the Global Burden of Disease 2019 estimate showing that globally in 2019, there were substantially more women than men living with dementia, with a female-to-male ratio of 1.69 (95% UI 1.64–1.73)^39^. For bipolar disorder, U.S. National Comorbidity Survey data indicate nearly equal prevalence—2.8% in women versus 2.9% in men—resulting in fractions of 0.491 and 0.509, respectively^40^. Dementia with Lewy bodies is markedly male-predominant: in Olmsted County, Minnesota, incidence was 4.8 per 100,000 person-years in men compared to 2.2 in women, yielding fractions of 0.314 and 0.686^41^. Frontotemporal dementia overall shows little sex bias, with pooled case series reporting near parity (373 males vs. 338 females), translating to fractions of 0.525 and 0.475^42^. Parkinson’s disease consistently demonstrates male predominance in North America, with men approximately 1.5 times more likely than women to develop the disease, corresponding to fractions of 0.600 and 0.400^43^. Schizophrenia also occurs more frequently in men, with meta-analyses estimating a male-to-female incidence rate ratio of ∼1.42:1, equivalent to 0.587 male and 0.413 female fractions^44^. Tauopathies as an aggregate entity are epidemiologically heterogeneous: progressive supranuclear palsy often shows modest male predominance, while corticobasal degeneration is closer to balanced; given the inconsistency, we reported a neutral 0.5/0.5 split as an aggregate reference^45^. Finally, vascular dementia appears more common in men in high-income cohorts, with estimates such as 5.6 vs. 3.2 cases per 1,000 person-years, corresponding to 0.636 male and 0.364 female fractions^46^.

### Curation of Demographic Data in Single-Cell Consortia Studies

#### Human Cell Atlas (HCA)

##### Data Source

We used publicly available datasets from the Human Cell Atlas (HCA) Bionetwork^1-3^ to assess ethnic disparities in single-cell genomics research. The analysis covered datasets from 14 HCA networks, including: adipose, breast, development, eye, gut, heart, immune, kidney, liver, musculoskeletal, nervous system, oral and craniofacial, pancreas, reproduction, and skin. Two networks, genetic diversity and organoid, did not have available datasets and were excluded from this analysis.

##### Dataset Selection and Metadata Extraction

For each network, we accessed all available datasets to review the associated metadata files, typically provided in Excel spreadsheet format. These metadata files were examined to identify clinical and demographic information about the donors, specifically focusing on age, gender, ethnicity, and disease status. If demographic metadata was present, we extracted and compiled the relevant information into a single comprehensive spreadsheet for each network.

In cases where ethnicity data was absent from the metadata, we manually reviewed the publications associated with the dataset. These publications were linked within each dataset’s page and often contained supplementary information. Ethnicity data was most frequently found in the Materials and Methods sections or the supplementary materials of these publications. We documented the presence or absence of metadata for each dataset and categorized the ethnicity reporting as follows:

1. Metadata Availability: We recorded whether the study included metadata with clinical and demographic data.
2. Ethnicity Reporting in Metadata: If ethnicity was present in the metadata, we marked it as reported.
3. Ethnicity Reporting in Publications: If no ethnicity data was found in the metadata, we documented whether it was available in the publication.

If ethnicity was provided in the metadata, the column asking about ethnicity data in publications was marked as “N/A.” In cases where the ethnicity data in the publication were incomplete or unclear (e.g., presented as proportions or graphs without clear detail), we flagged this limitation with an asterisk.

##### Standardization of Ancestry and Ethnicity Categories

Given the variability in how ethnicity data were reported across projects (e.g., “Asian,” “East Asian,” or “Korean”), we standardized the ethnicity categories to allow for consistent comparisons across datasets. The following broad categories were used:

- European
- African
- East Asian
- South Asian
- Asian: For cases where further specification was not provided.
- American: Including Hispanic/Latino and Native American individuals.
- Other: Encompassing other ethnic groups not covered by the categories above and individuals of mixed ethnic backgrounds.
- Unknown

All Asian categories were eventually merged in analyses due to the constraint from some studies. All more granular data were also preserved in the data available for download in this manuscript.

Donors from other species (e.g., Mus musculus) and cell lines, including human-induced pluripotent stem cells (iPSCs), were excluded from the dataset.

### HTAN (Human Tumor Atlas Network)

We analyzed HTAN (Human Tumor Atlas Network)^4,5^ data by filtering for datasets with single cell RNA sequencing data (scRNA-seq). For each HTAN dataset, we accessed the associated metadata files and reviewed sample-level attributes including primary tissue of origin, tumor type, anatomical location, and contributing research center. To ensure consistency across datasets, we created a standardized labeling system for tissue origin and tumor site, modeled after the category structure used on the HTAN homepage. Where metadata fields were ambiguous or inconsistent, we manually reconciled them using information from the associated publications and supplementary materials. Each sample was then re-annotated using this uniform classification framework, allowing for improved comparison across atlases. Given the limited number of samples in certain cancer types, all common hematologic malignancies to a single Blood Tumors category, and bone / bone marrow / joint sites are merged to a Bones category. The curated metadata was compiled into a master file that enables streamlined analysis of tissue-specific patterns and site-based tumor characteristics.

To classify ancestry groups, metadata for scRNA-seq from HTAN were extracted and analyzed for racial/ethnic differences to be processed. For the purpose of this study, individuals reported as Hispanic/Latino were considered as Latino, individuals reported White but non-Latino or missing ethnicity were considered as European, individuals reported as African or African American were considered as African, individuals reported to be East Asian, South Asian, or EAS_SAS were considered Asian, individuals reported to be other values were considered as Other, and Unknown or missing data were considered as Unknown.

### PsychAD Consortium

We downloaded the metadata from the Supplementary Table 1 of PsychAD’s latest data release and landmark manuscript^6^ as of May 2025. In this metadata, ancestries of AFR is mapped to African, AMR to Latino, EUR to Europeans, and EAS/SAS/EAS_SAS to Asians. To identify each sample’s disease status, we simply used the coded variables where an individual could have more than one diagnosis, and all others without any diagnosis is considered as controls.

## Tables

**Table 2. Key design considerations for ensuring fair representation in single-cell studies**

## Supplementary Tables

**Supplementary Table 1. Human Cell Atlas (HCA) dataset ancestry and sex composition, shown in counts and percentages within each tissue type**

**Supplementary Table 2. Robustness analyses of ancestry representation in the Human Cell Atlas (HCA) dataset**

**Supplementary Table 3. Human Tumor Atlas Network (HTAN) ancestry and sex composition, shown in counts and percentages within each cancer type**

**Supplementary Figure 4. PsychAD dataset ancestry and sex composition**,, **shown in counts and percentages within each disease type**

## DATA AND SOFTWARE AVAILABILITY

### Data Availability

The curated metadata datasets for this study, including HCA, HTAN, and PsychAD and their reference datasets, are deposited in https://zenodo.org/records/17161565

### Code Availability

The source code to reproduce all analyses can be found at: https://github.com/Huang-lab/scOmics_equity under the open source MIT license.

## ACKNOWLEDGMENTS

The authors wish to acknowledge the Dataworks! Prize hosted by the Federation of American Societies for Experimental Biology (FASEB) and the National Institutes of Health (NIH) for inspiring us to conduct this study. The study was based on volunteer efforts; the publication is funded by NIH NIA/NHGRI UG3AG105083, NIGMS R35GM138113 and 2R35GM138113 to K.H.

## COMPETING FINANCIAL INTERESTS

K.H. is a co-founder and board member of a 501(c)(3) not-for-profit Open Box Science (unpaid) and Kaimen Inc., both of which were unrelated to this study. All other authors declare no competing interests.

## CONTRIBUTIONS

K.H. conceived the research and designed the analyses. C.Y. curated the HCA data, A.S. curated the HTAN data, and K.S. conducted the exploratory analyses. K.H. curated the PsychAD data and reference data. K.H. conducted all final analyses and supervised the study. K.H. wrote the manuscript; all authors read, edited, and approved the manuscript.

## Notes

https://zenodo.org/records/17161565

### Summary of Updates

1. Deepening the Biological Rationale and Single-Cell Specificity We added a new section in the Introduction articulating why ancestry matters for single-cell biology on its own terms, moving beyond the GWAS analogy. We now cite single-cell studies demonstrating ancestry-differential gene expression in immune cell subsets (Randolph et al., Science 2021; N et al., Cell 2016; Aquino et al., Nature 2023), concrete pharmacogenomic examples (CYP2D6 population variation), and the risk of missing cell subtypes in homogeneous atlases. We introduce the "pancellular reference" concept, framing diverse atlases as a scientific prerequisite rather than solely a fairness imperative. 2. Moving from Theoretical AI Risk to Empirical Evidence We reframed claims about AI foundation models as structural risk supported by a training-data audit. We explicitly identify that scGPT, Geneformer, and scFoundation draw on HCA/HTAN data, note the absence of controlled ancestry-stratified benchmarks, and add the point that inequitable access to computational infrastructure (GPUs, AI tools) constitutes a secondary layer of disparity. 3. Methodological Rigor, Metadata Gaps, and Global Scope We added partial-identification sensitivity analyses for HCA, reporting bounded ancestry ranges under plausible missing-data mechanisms (Supplementary Table 2). We now explicitly discuss study-purpose-driven sampling, acknowledge that non-consortium datasets, and describe our metadata curation process in addition to Methods. We discuss ongoing efforts toward metadata standardization (cellxgene, EBI, HCA Diversity Task Force) and genetically inferred ancestry from scRNA-seq data. 4. Addressing Structural, Political, and Economic Barriers A new discussion section addresses the intersecting forces driving demographic imbalance: convenience-based study design, Western-centered funding structures, national data-governance restrictions, high technology costs, and the need for respectful community engagement, including Indigenous data sovereignty frameworks. 5. Actionable, Field-Specific Guidance Table 2 has been completely rewritten as a 16-question, 5-domain checklist spanning the full study lifecycle, from procurement and recruitment through pre-analytical considerations, equitable multiplexing/batch design, metadata ontology standards, consent and data sovereignty, to AI model reporting requirements. We also survey the growing ecosystem of diversity initiatives (HCA Equity Working Group, CZI Ancestry Networks, regional atlas projects) to ground our recommendations in what the community is already doing.

