## Supplementary figures and images for "Are Different Populations Fairly Represented in Single-Cell Omic Atlases?"

### Supplementary Figures

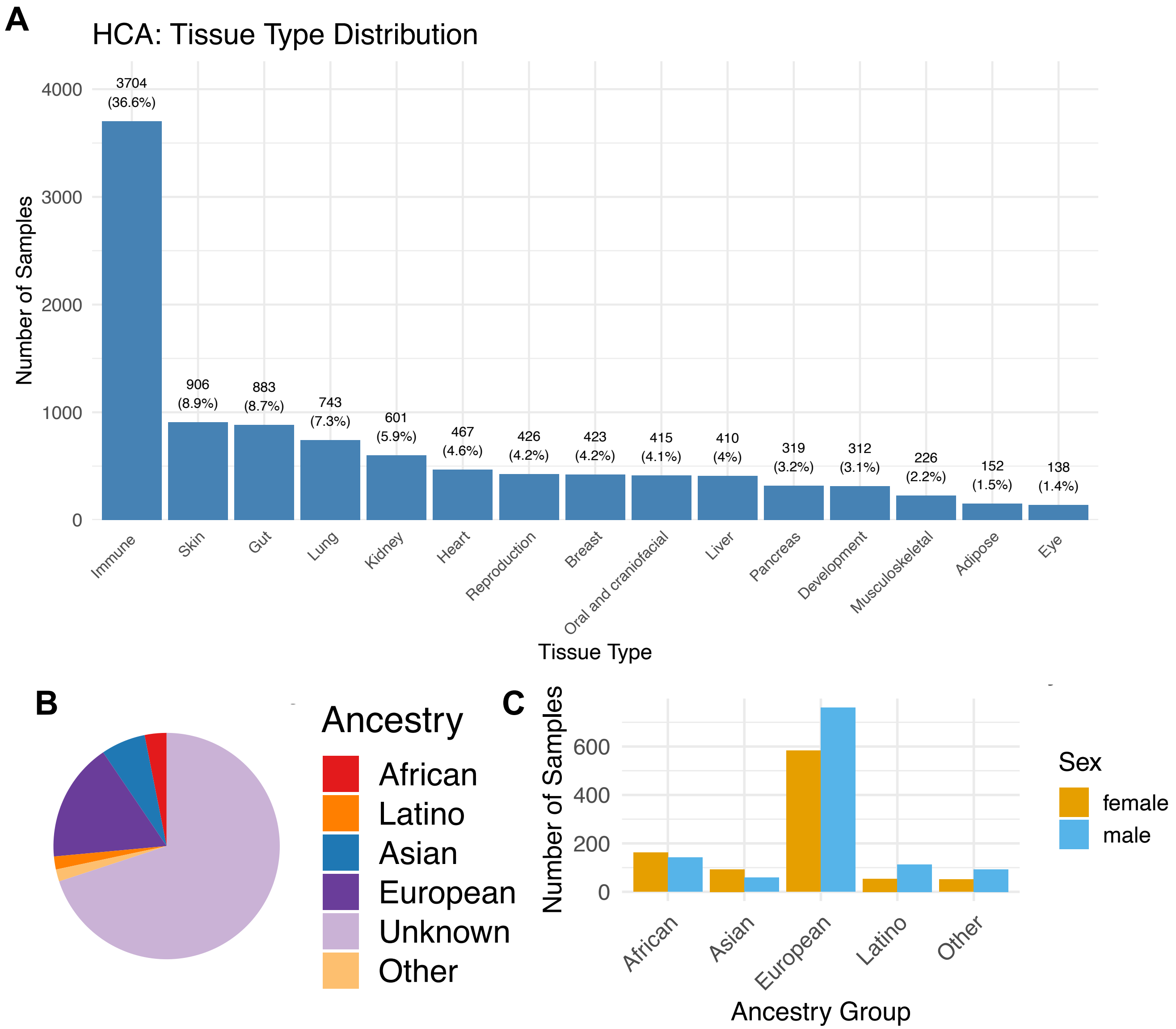

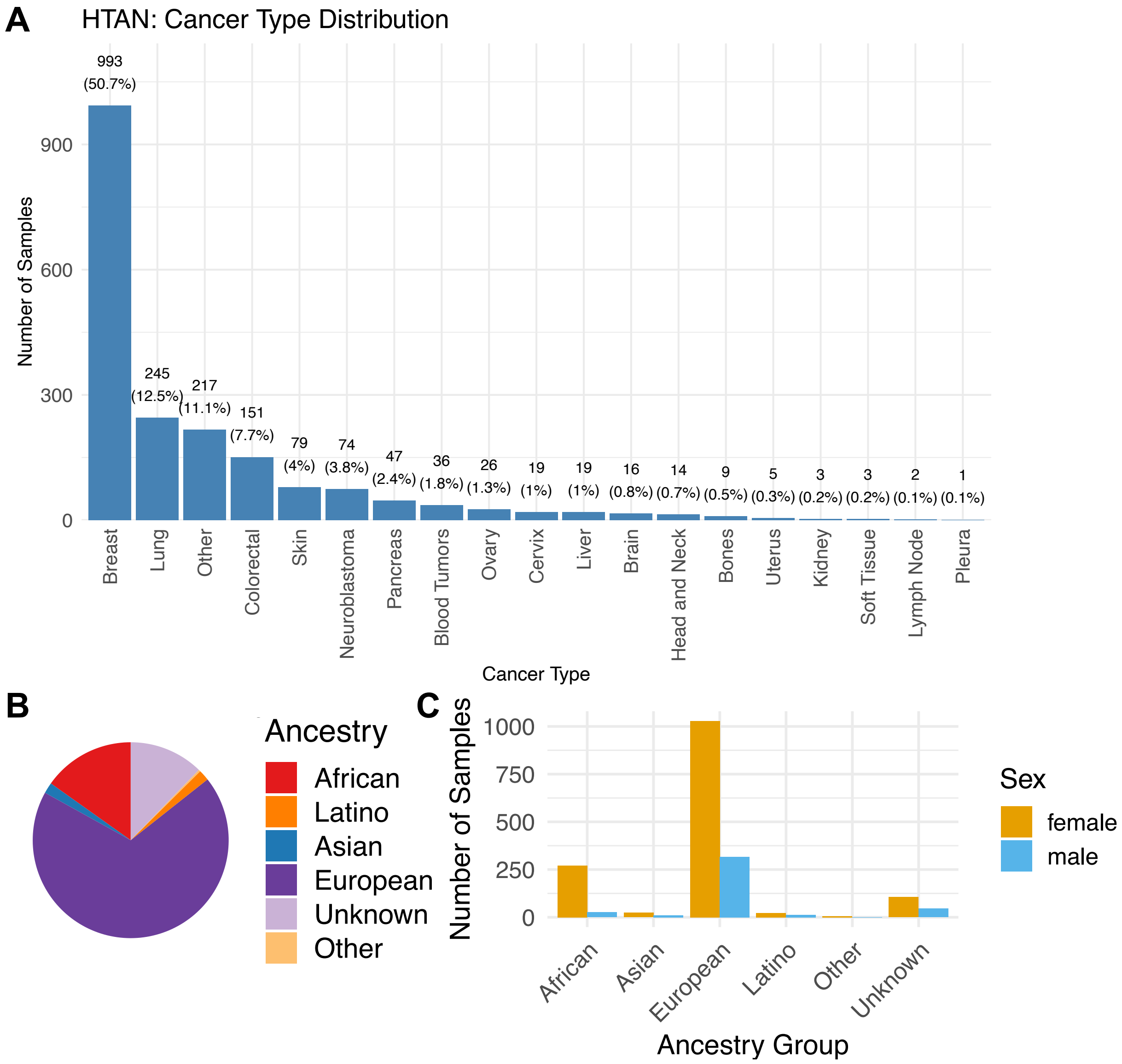

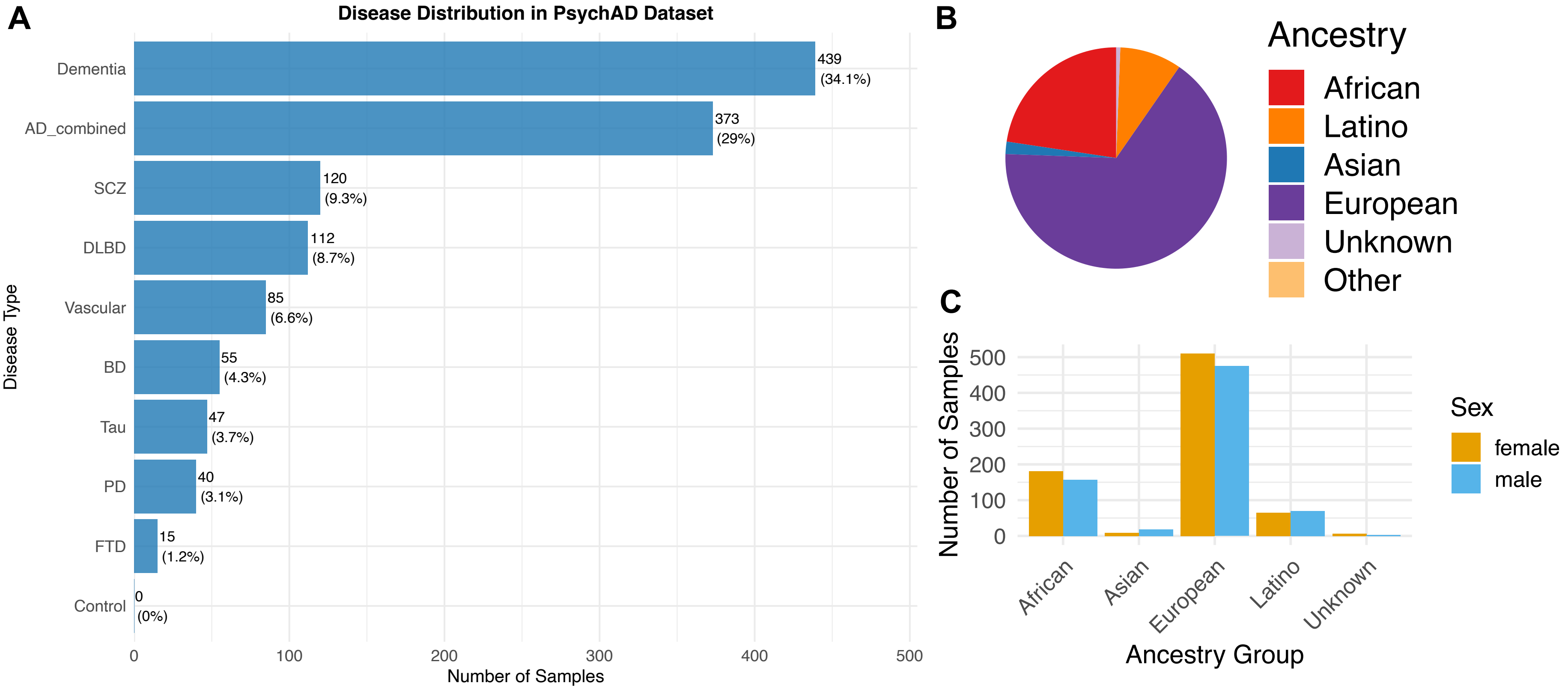
